# Integrating trajectory inference and gene regulatory network analysis to resolve transcriptional programs of T cell state transitions in the tumor microenvironment

**DOI:** 10.64898/2026.05.12.724558

**Authors:** Roger Casals-Franch, Lara Nonell, Jordi Villà-Freixa, Adrian Lopez Garcia de Lomana

## Abstract

Reconstructing dynamic immune cell state transitions from single-cell transcriptomic data requires coordinated analytical strategies that capture both phenotypic progression and underlying regulatory programs. This protocol describes a step-by-step computational workflow for analyzing human tumor-infiltrating T cells using the sequential application of dimensionality reduction, pseudotime trajectory inference, regulon activity analysis, and transcription factor–transcription factor network reconstruction. The workflow outlines data preprocessing and quality control, trajectory rooting and parameter selection, branch-specific differential analysis, and the integration of regulon inference to contextualize transcriptional programs along inferred trajectories. Regulon-based TF–TF network reconstruction is used as a downstream interpretive layer to identify regulatory modules associated with distinct cell-state transitions. Publicly available at GitHub repository https://github.com/rogercasalsfr/immuno-trajectory-grn-integrative-workflow, this protocol emphasizes practical considerations including parameter sensitivity, trajectory robustness, and consistency between phenotypic and regulatory outputs. The protocol supports reproducible analysis and interpretation of immune cell dynamics in human tumor microenvironment studies using single-cell RNA sequencing data.

## Introduction

The tumor microenvironment (TME) is a complex and dynamic ecosystem in which immune cells continuously adapt their functional states in response to tumor-derived signals and therapeutic interventions [1,2]. Among these, T cells are central mediators of anti-tumor immunity, and their transcriptional heterogeneity and plasticity fundamentally shape the outcome of cancer immunotherapy [3]. Immune checkpoint blockade (ICB) therapies, particularly those targeting the PD-1/PD-L1 axis, have demonstrated clinical efficacy across multiple cancer types by restoring T-cell function in the TME [4]. However, the molecular mechanisms governing T-cell state transitions in response to therapy remain incompletely understood, and a deeper characterization of these regulatory processes is needed to improve therapeutic design and patient stratification.

Single-cell RNA sequencing (scRNA-seq) has transformed our capacity to study immune cell biology by enabling the simultaneous profiling of thousands of individual cells at transcriptomic resolution [5]. A particularly powerful application of scRNA-seq is the inference of differentiation trajectories, which reconstructs the dynamic progression of cell states from transcriptomic snapshots. Trajectory inference methods, such as Monocle 3 [6], model cell-state transitions by embedding cells in a low-dimensional space and fitting a principal graph that captures the topology of differentiation. This approach assigns each cell a pseudotime value — a continuous, time-like coordinate that reflects its progression along a given differentiation path — without requiring longitudinal sampling. When applied to tumor-infiltrating T cells, trajectory inference can reveal how therapy reshapes differentiation dynamics, identify branch points corresponding to fate decisions, and highlight the gene expression changes that accompany each transition.

Trajectory inference alone, however, captures phenotypic progression without revealing the regulatory mechanisms that drive it. Gene regulatory network (GRN) inference addresses this limitation by identifying coherent programs of transcription factor (TF) activity and their target gene sets, termed regulons. Tools such as pySCENIC [7] implement a three-step workflow that combines co-expression analysis, cis-regulatory motif enrichment, and single-cell activity scoring to reconstruct regulon activity at single-cell resolution. By integrating regulon activity with trajectory analysis, it becomes possible not only to characterize what cells are doing at each state, but also to identify which transcriptional programs are driving those states and how they are rewired by external perturbations such as immunotherapy.

This protocol describes a step-by-step computational workflow that integrates these complementary approaches into a coherent analytical framework for the study of T-cell dynamics in the human TME [8]. Starting from a publicly available scRNA-seq dataset of tumor-infiltrating T cells from squamous cell carcinoma patients sampled before and after anti-PD-1 therapy [9], the workflow guides the reader through data preprocessing and quality control, pseudotime trajectory inference using a variety of tools (Monocle 3 [6], PAGA [10] and Slingshot [11]), branch-specific differential expression and pathway over-representation analysis using Reactome [12], GRN inference using pySCENIC [7], and downstream visualization of TF–TF interaction networks in Cytoscape [13]. While the protocol is demonstrated on this specific dataset, the analytical strategy is broadly applicable to other scRNA-seq datasets of immune cells in which discrete functional states can be annotated and compared across conditions.

Several practical considerations are central to the successful application of this workflow. First, trajectory inference is sensitive to the quality of cell-type annotation: the assignment of a trajectory root and the biological interpretation of inferred branches both depend on prior knowledge of the expected differentiation hierarchy. This protocol therefore recommends using established canonical marker genes to guide annotation and branch labeling, and includes guidance on how to select appropriate root populations under different experimental conditions. Second, GRN inference using pySCENIC is computationally intensive, particularly the motif enrichment step, and is recommended to be run in a containerized environment using the provided Docker image to ensure software compatibility and reproducibility. Third, the integration of trajectory and regulon outputs requires careful alignment of cell identifiers and metadata between R-based and Python-based analysis environments; this protocol provides explicit guidance on data export and import steps to facilitate this integration.

Together, this workflow enables the reconstruction of differentiation trajectories, the identification of branch-specific gene expression programs, and the characterization of the transcriptional regulatory networks that govern T-cell state transitions in the context of immunotherapy. The protocol is designed to support researchers with familiarity in basic command-line usage, and all analysis scripts are publicly available at GitHub repository https://github.com/rogercasalsfr/immuno-trajectory-grn-integrative-workflow to facilitate reproducibility and adaptation to new datasets.

## 2. Materials

### 2.1. Software and computational environment

1. Linux workstation, server, or high performance computing (HPC) environment with Apptainer or SingularityCE [14] for containerized execution.
2. The working repository for this workflow, https://github.com/rogercasalsfr/immuno-trajectory-grn-integrative-workflow. This repository includes Apptainer/Singularity definition file env/pstime.def, conda environment files env/environment.yml and env/pstime.yml and Python notebooks workflow/protocol.ipynb
3. Python 3.14.3 environment containing at least the core packages recorded in env/environment.yml, including anndata 0.12.6, scanpy 1.12, decoupler-py 2.1.4, pandas 2.3.3, numpy 1.26.4, scipy 1.17.1, scikit-learn 1.8.0, umap-learn 0.5.11, py-monocle 0.1.1, pyslingshot 0.2.0, networkx 3.6.1, matplotlib 3.10.8, and jupyterlab 4.5.5.
4. Optional annotation software such as celltypist [15] when manual annotations are not supplied by the source dataset, as in this protocol.
5. pySCENIC 0.12.1 [7] available as a Docker container image for upstream regulon inference when raw-expression GRN reconstruction is required.
6. Cytoscape [13] or equivalent graph visualization software for TF-TF network display.

### 2.2. Hardware requirements

1. A personal workstation with at least 16 GB RAM and 8 CPU cores is sufficient for a reduced tutorial-scale analysis.
2. Larger datasets and the pySCENIC motif-pruning step benefit larger computational environments, as in a cluster.
3. For full pySCENIC processing, plan for high-memory nodes; the regulon inference step is treated as an external analysis in this protocol and may require up to 128 GB RAM depending on dataset size

### 2.3. Input data & metadata

1. GEO accession GSE123813 [16], including the raw count matrix and cell-level metadata, including cell type annotations.
2. Cell metadata containing at least cell barcodes, patient identifiers, treatment labels, and, when available, as in this protocol, source-study cell-type annotations (see **Note #1**).

### 2.4. Reference files for gene regulatory network analysis

1. In this protocol, for computational feasibility, we provide the pySCENIC outputs to seamlessly integrate with the trajectory inference results. Precomputed pySCENIC GRN files provided are `expr_mat.adjacencies.tsv` and `regulons.csv`, all available under `workflow/GRN_inference` folder in the GitHub repository (see **Note #2**).

## 3. Methods

### 3.1. Create a reproducible analysis environment

1. Clone or download the working repository https://github.com/rogercasalsfr/immuno-trajectory-grn-integrative-workflow to a Linux-accessible working directory.
2. Build the container image from the provided recipe when reproducible execution is required. A typical command is apptainer build `--fakeroot pstime.sif env/pstime.def` (see Note #3).
3. If a container is not used, recreate the environment from env/environment.yml, making sure that the package versions remain synchronized with the notebook used for the analysis.
4. Launch JupyterLab inside the environment and open workflow/protocol.ipynb as the working notebook (see Note #4).

### 3.2. Download and organize the example dataset

1. Create a dedicated data directory, for example data/GSE123813.
2. Download the metadata and count matrix (cells as rows and genes as columns) files associated with GEO accession GSE123813.
3. Keep the original file names and document the download date if the protocol is being reused as a teaching or benchmarking resource.
4. Store the raw input files separately from derived objects such as filtered matrices, figures, and pseudotime outputs.

### 3.3. Load the count matrix and metadata into an AnnData object in the Python notebook environment

1. Read the count matrix and metadata table into the Python notebook.
2. Transpose the count matrix if needed so that cells are rows and genes are columns before creating the AnnData object.
3. Align the metadata to the expression matrix by cell barcode and confirm that the order of cells is identical in both objects (see Note #5).
4. Record the initial dimensions of the dataset and inspect whether duplicated barcodes or missing metadata entries are present.

### 3.4. Inspect the dataset and define the data subset depending on analysis scope

1. Review the number of cells, number of genes, available metadata fields, treatment labels, and patient representation.
2. Determine whether the study question should be addressed jointly across all T cells or separately within biologically coherent compartments such as CD4^+^ and CD8^+^ cells.
3. Inspect whether pre-treatment and post-treatment cells occupy overlapping manifolds or whether they form strongly separated state spaces.
4. If treatment conditions are topologically disconnected or dominated by condition-specific cell states, plan separate trajectory analyses and compare the results afterward rather than forcing all cells into a single pseudotime. Large treatment-induced composition shifts are often better handled with separate trajectories and a later comparison.

### 3.5. Select the cells of interest and confirm cell-state annotations

1. Subset the dataset to the T-cell compartment relevant to the biological question.
2. In this workflow, analyses on CD4^+^ and CD8^+^ cells are treated separately as they represent different differentiation programs. We will proceed with the CD4^+^ cells subset.
3. Use prior author-provided annotations when available, and refine them eventually with canonical marker genes.
4. If annotations are absent or insufficiently resolved, use an automated annotation framework such as CellTypist [15] as a starting point, and always verify the resulting labels against canonical marker gene expression before proceeding.
5. Confirm that the selected cell population is sufficiently large for downstream analyses. As a practical guideline, aim for at least 200 cells per condition or branch. Very small groups can destabilize branch-specific differential expression and regulon comparisons.
6. Confirm that expected markers support the chosen labels. For example, in this workflow, markers reported in Yost et al such as *CCR7* for naive-like cells, *FOXP3* for Tregs, and *CXCL13* for Tfh states help validate annotation quality (see Note #6).

### 3.6. Perform quality control and normalization

1. Filter genes detected in very few cells, for example fewer than 5 cells.
2. Remove low-quality cells using dataset-specific thresholds for total UMI counts, number of detected genes, and mitochondrial transcript percentage.
3. Normalize counts per cell to a fixed library size, apply log transformation, identify highly variable genes, scale the matrix, and compute principal components.
4. Inspect quality control variable distributions before fixing thresholds and document any deviations from the default settings (see **Note #7**).

### 3.7. Correct patient effects, cluster the cells, and compute a UMAP embedding

1. Compute PCA on the filtered and normalized matrix.
2. If patient identity is a dominant source of variation, apply Harmony [17] to the PCA coordinates using patient ID as the batch variable.
3. Build the nearest-neighbor graph on the corrected latent space, calculate Leiden clusters, and compute a UMAP embedding for visualization.
4. Plot the embedding colored by cell type, patient, and treatment condition to confirm that the main structure reflects biology rather than patient-specific artifacts (**Fig. 1**).
5. Tune clustering resolution and neighborhood parameters according to the biological granularity of interest, and document the values used in the final analysis (see **Note #8**).

**Figure 1.**
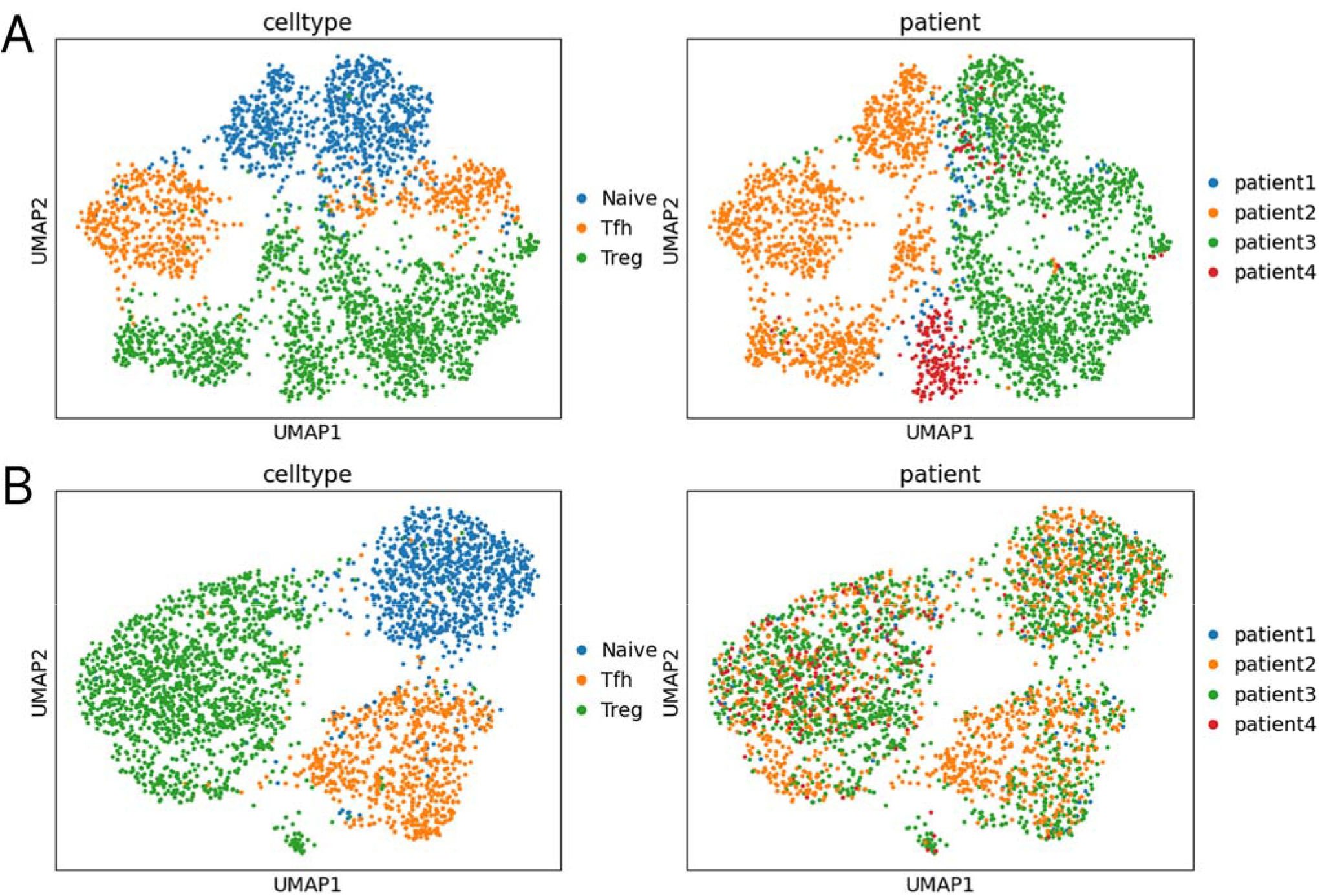
UMAP embeddings of CD4^+^ tumor-infiltrating T cells before and after batch effect correction. Each panel shows the same UMAP projection colored by cell type (left) and patient identity (right). **A**. Before correction, batch effects driven by patient identity are a dominant source of variation, with cells from individual donors forming largely separate clusters. **B**. After Harmony-based batch effect correction, cells from different donors are well intermixed while cell-type structure is preserved, indicating successful removal of patient-driven technical variation without loss of biological signal.

### 3.8. Choose a biologically plausible trajectory root

1. Select the trajectory root using prior biological knowledge, marker expression, and cluster identity rather than visual position alone.
2. In the example workflow, naive-like T cells are used as the default root because they represent the least differentiated state within the analyzed continuum.
3. When a truly naive population is absent, choose the earliest biologically plausible precursor state, such as a memory-like or stem-like compartment, and justify the choice explicitly (see **Note #9**).
4. Save the root definition together with the final metadata so that the pseudotime orientation can be reproduced exactly.

### 3.9. Infer trajectories with independent methods for robustness assessment

#### 3.9.1. PAGA

1. Run PAGA [10] on the annotated states or on unsupervised clusters to obtain a coarse-grained view of connectivity.
2. Use the PAGA graph, which represents the connectivity between cell clusters as a topology-preserving abstraction of the data manifold, to determine whether the selected cell subset is fully connected or contains isolated populations that should be excluded from trajectory analysis (**Fig. 2A**).
3. Remove clearly disconnected populations if they do not belong to the biological process under study.

**Figure 2.**
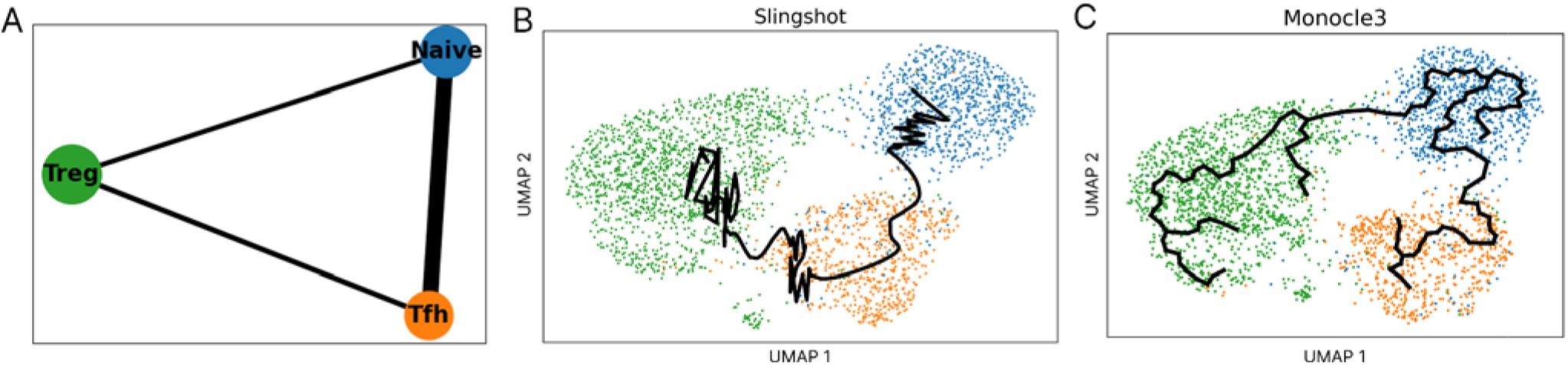
Trajectory inference for CD4^+^ tumor-infiltrating T cells across three independent methods. **A**. PAGA connectivity graph showing inferred relationships between naive, Tfh, and Treg clusters; edge weights reflect connectivity strength. **B**. Slingshot lineage curves on UMAP colored by cell type. Slingshot recovers a Tfh-to-Treg transition unsupported by the other methods or canonical CD4^+^ T cell biology, and misses the expected naive-to-Treg path. **C**. Monocle 3 pseudotime trajectory on the same UMAP, rooted at naive cells, recovering both naive-to-Treg and naive-to-Tfh paths. Agreement between PAGA and Monocle 3 supports the inferred topology; the Slingshot discrepancies highlight the value of cross-validating trajectory inference across methods.

**Figure 3.**
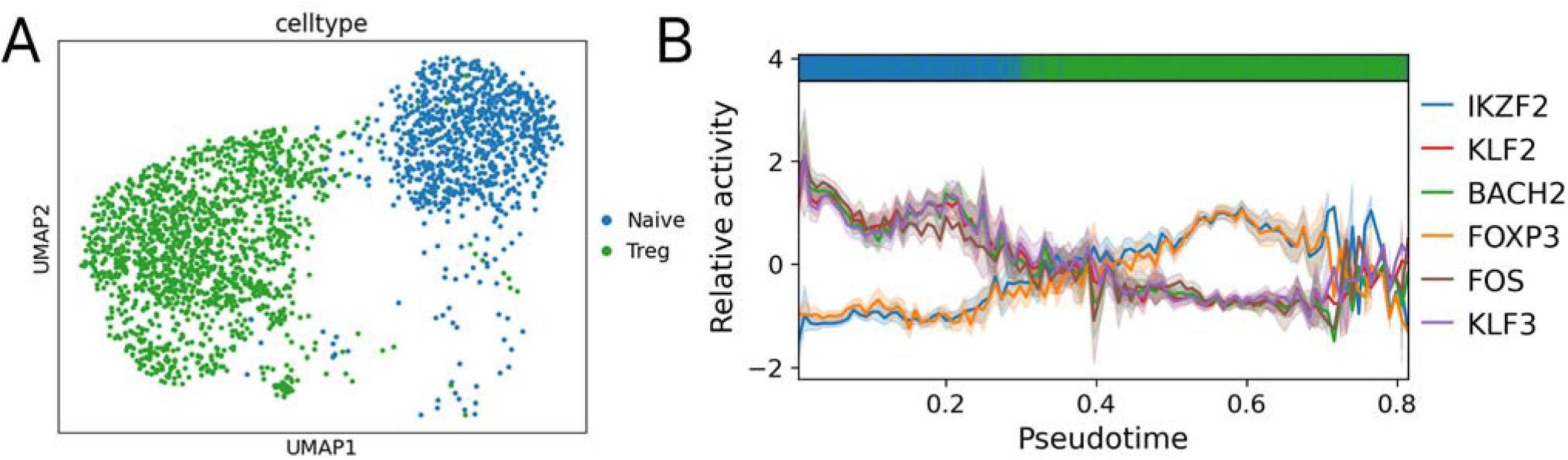
Transcription factor activity along the naive-to-Treg differentiation trajectory in CD4^+^ tumor-infiltrating T cells. **A**. UMAP embedding of strictly naive and Treg cell populations colored by cell type. **B**. Smoothed regulon activity scores for the five transcription factors most significantly associated with the naive-to-Treg transition (KLF2, BACH2, IKZF2, FOXP3, and FOS) plotted along Monocle 3 pseudotime. Cell type annotations are indicated along the x-axis. As expected, KLF2 activity decreases along the trajectory while FOXP3 activity increases, consistent with the transition from a naive to a regulatory T cell state.

**Figure 4.**
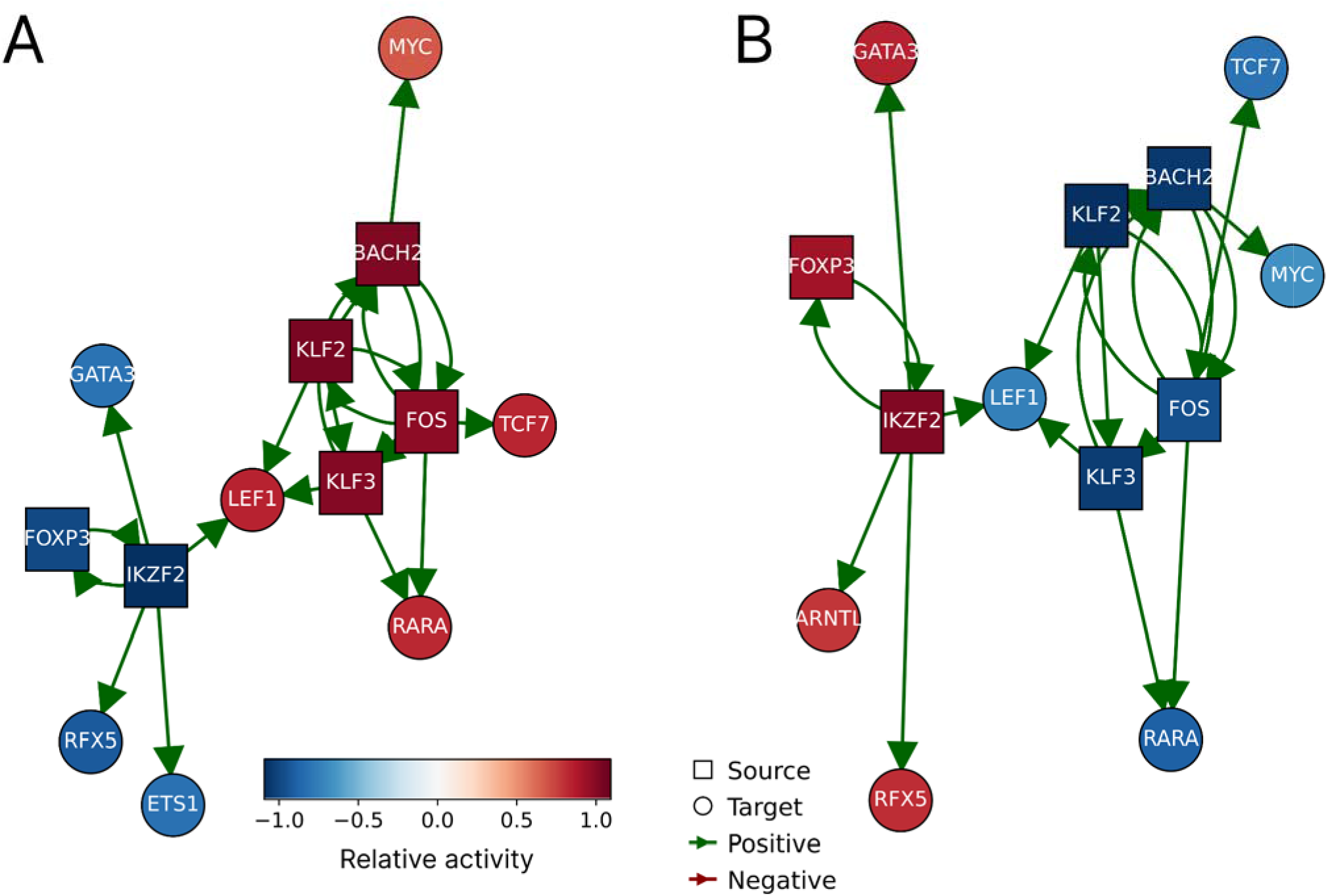
TF-TF regulatory subnetwork for the naive and Treg programs in CD4^+^ tumor-infiltrating T cells. **A**. Network activity in the naive population. **B**. Network activity in the Treg population. In both panels, nodes represent regulons (labelled by their controlling TF) and edges indicate regulatory interactions derived from the inference step (using scenic) towards any of the source nodes. Node color reflects regulon relative activity across cells, with red indicating positive and blue indicating negative activity. No negative regulatory interactions are found in this network.

#### 3.9.2. Monocle 3

1. Learn a principal graph on the selected embedding and assign pseudotime values from the chosen root cells using Monocle 3 [6].
2. Visualize pseudotime and the principal graph on the embedding (**Figure 2B**).
3. Inspect sensitivity to neighborhood construction, embedding parameters, and partitioning behavior before fixing the final trajectory.

#### 3.9.3. Slingshot

1. Run Slingshot [11] using cluster labels and a biologically justified starting cluster (**Fig. 2C**).
2. Compare the inferred lineage structure with the Monocle 3 result.
3. Use Slingshot as a robustness analysis to determine whether broad lineage structure is stable across trajectory methods.

#### 3.9.4. Compare outputs across methods

1. Accept a trajectory only when topology, marker trends, and biological interpretation agree across PAGA, Monocle 3, Slingshot, and relevant literature.
2. If the methods disagree strongly, revisit the selected cell subset, annotation strategy, batch correction, or root definition before proceeding (see **Note #10**).

### 3.10. Estimate transcription factor activity along pseudotime with decoupler

1. Retrieve a human TF-target prior network, for example CollecTRI [18], using the decoupler-py interface [19].
2. Score TF activity on the single-cell object by applying the univariate linear model (ULM) using `dc.run_ulm()`, with the CollecTRI prior network as input, and store the resulting TF activity matrix in the AnnData object for downstream visualization and ranking (see **Note #11**).
3. Rank TFs according to their association with pseudotime, for example by distance correlation against the pseudotime order.
4. Visualize the top regulators on the UMAP and as binned trends along pseudotime.

### 3.11. Identify genes associated with pseudotime progression and branch divergence

1. Use a graph-aware statistic such as Moran’s I as implemented in Monocle 3 to identify genes that vary smoothly along the inferred manifold.
2. Plot the top pseudotime-associated genes on the embedding to confirm that their behavior is consistent with the proposed trajectory.
3. Perform additional differential expression analyses between branches, terminal states, treatment conditions, or manually defined trajectory segments as needed.
4. Keep the statistical framework consistent across comparisons and state clearly whether the analysis is pseudotime-driven, branch-driven, or condition-driven.

### 3.12. Perform pathway enrichment analysis

1. Export significant pseudotime-associated genes for pathway over-representation analysis using resources such as Reactome [12] or Enrichr [20].
2. Complement gene-list enrichment with single-cell pathway scoring by applying the ULM (`dc.run_ulm()`) from decoupler-py to the MSigDB Hallmark gene sets [21], accessed directly via `dc.op.hallmark()`, to quantify pathway activity at single-cell resolution across cell states and pseudotime intervals.
3. Compare pathway behavior across cell types, treatment conditions, and pseudotime intervals rather than interpreting pathway scores in isolation.

### 3.13. Integrate precomputed pySCENIC outputs for regulatory network analysis

1. When full GRN inference is required, execute the standalone pySCENIC pipeline using the standard three-step workflow: co-expression module detection (grn), cis-regulatory motif pruning to refine TF-target interactions (ctx), and regulon activity scoring (aucell). We provide this pipeline as a notebook `workflow/GRN_inference/GRN_inference.ipynb`. Alternatively, in this protocol, we provide the resulting outputs `expr_mat.adjacencies.tsv and regulons.csv`, already available in the `workflow/GRN_inference` folder (see **Note #12**).
2. Import the adjacency matrix and regulon definitions, keep the high-confidence TF-target edges, i.e., top 10%, and prune the network to interactions supported both by co-expression and motif-backed regulon enrichment.

### 3.14. Identify branch-associated regulatory programs

1. Subset the trajectory to the branch of interest, for example the naive-to-Treg arm.
2. Recompute GRN-based TF activity on the subset and rank the regulators by association with branch-specific pseudotime.
3. Compare branch-associated TFs with canonical markers, pathway results, and global pseudotime trends to prioritize candidate regulators that may drive lineage divergence.

### 3.15. Construct a TF-TF network and visualize candidate regulators of cell state transition

1. Filter the pySCENIC adjacency matrix to retain only edges where both source and target are known transcription factors, using a curated TF reference list such as Lambert et al. [22] downloaded from source, producing a TF-TF subnetwork (`tf_grn`).
2. Compute z-scored TF activity per cell, grouping cells by the median of annotated cell type (`df_naive, df_treg`).
3. Visualize the subnetwork with `dc.pl.network()`, passing tf_grn as net, and the median z-scored regulon activity scores as score. Source node color reflects regulon activity score; target node color reflects median target regulon activity score.
4. Compare networks across cell states to identify regulatory rewiring associated with the transition of interest.
5. Prioritize TFs that show both high regulon activity along pseudotime and high connectivity within the subnetwork (see **Note #13**).

## 4. Notes

1. User-provided single-cell RNA sequencing data in an equivalent genes-by-cells format may also be used if matching metadata are available.
2. The precomputed pySCENIC output files can be reproduced using the human transcription factor list for pySCENIC (allTFs_hg38.txt), available from the aertslab cisTarget resources server (https://resources.aertslab.org/cistarget/tf_lists/allTFs_hg38.txt). This list is derived from the curated inventory of human transcription factors compiled by Lambert et al. [22]. Additionally, human hg38 cisTarget ranking databases in Feather v2 format and a motif-to-TF annotation file, available from https://resources.aertslab.org/cistarget/databases/, are required for the context pruning step of pySCENIC [7].
3. Please refer to https://apptainer.org/docs/user/latest/ for guidance on building containers.
4. Use a fixed environment for the full workflow. Small differences in package versions can change the nearest-neighbor graph, clustering, UMAP layout, and downstream activity scores. Archive the final environment definition or container image together with the notebook outputs so that the protocol remains reproducible on HPC systems and across future software versions.
5. Do not continue until cell identifiers match exactly between the metadata and the expression matrix. Misaligned metadata will corrupt downstream biological interpretations.
6. Annotation quality is one of the main determinants of whether trajectory inference will be biologically interpretable. A poor annotation step propagates directly into root selection, branch labeling, and regulatory interpretation.
7. The thresholds used in the example notebook are dataset-specific examples rather than universal defaults. Overly permissive filtering retains damaged cells, whereas overly stringent filtering can remove rare transitional states that are essential for trajectory analysis.
8. The choice of Leiden resolution, neighborhood size, batch-correction strategy, and UMAP parameters can all change the apparent topology. Treat these settings as sensitivity-analysis parameters rather than fixed truths. Additionally, UMAP is useful for visualization, but trajectory-learning methods can be sensitive to embedding choices. Important conclusions should be cross-checked against PCA and batch-corrected latent spaces rather than accepted solely on the basis of a visually appealing embedding.
9. Pseudotime is inferred, not measured. It provides a relative ordering of cells within the selected manifold rather than an absolute temporal axis. Root choice determines pseudotime direction, as an implausible root can invert biological interpretation and change the ranking of genes, pathways, and regulators associated with progression.
10. Not every bifurcation in the inferred graph corresponds to a true biological fate decision. Branch points must be supported by marker gene behavior, pathway enrichment, and consistency with published literature before being interpreted as biologically meaningful transitions.
11. TF activity scores inferred with decoupler summarize coordinated target-gene behavior and should not be interpreted as direct measurements of TF transcript abundance. Similarly, pySCENIC-derived regulons and decoupler-based activity scores are hypothesis-generating outputs, as they prioritize candidate regulatory mechanisms for follow-up investigation and should not be interpreted as proof of direct causality.
12. pySCENIC motif pruning is memory-intensive and is best run in a containerized HPC setting. Plan for up to 128 GB RAM depending on dataset size. Retain the exact database versions and command lines used for the upstream GRN run, as these are required for full reproducibility.
13. If a key transcription factor shows weak or undetectable mRNA expression in the dataset, do not interpret its absence as biological inactivity. Inspect target-gene behavior and regulon activity scores before concluding that the regulator is inactive, as mRNA abundance and regulatory activity are not equivalent.

## Acknowledgements

The authors acknowledge the computational resources provided by the Mímir cluster of the Icelandic Research e-Infrastructure (IREI), funded by the Icelandic Centre of Research infrastructure fund. This work was supported by the Investigo Programme, funded by the European Union – NextGenerationEU and the Generalitat de Catalunya (supporting RCF).

